# Modeling the growth and decline of pathogen effective population size provides insight into epidemic dynamics and drivers of antimicrobial resistance

**DOI:** 10.1101/210054

**Authors:** Erik M. Volz, Xavier Didelot

**Author notes:** **Corresponding author:** Erik Volz, Department of Infectious Disease Epidemiology, Imperial College London, Norfolk Place, W2 1PG, United Kingdom.

## Abstract

Non-parametric population genetic modeling provides a simple and flexible approach for studying demographic history and epidemic dynamics using pathogen sequence data. Existing Bayesian approaches are premised on stationary stochastic processes which may provide an unrealistic prior for epidemic histories which feature extended period of exponential growth or decline. We show that non-parametric models defined in terms of the growth rate of the effective population size can provide a more realistic prior for epidemic history. We propose a non-parametric autoregressive model on the growth rate as a prior for effective population size, which corresponds to the dynamics expected under many epidemic situations. We demonstrate the use of this model within a Bayesian phylodynamic inference framework. Our method correctly reconstructs trends of epidemic growth and decline from pathogen genealogies even when genealogical data is sparse and conventional skyline estimators erroneously predict stable population size. We also propose a regression approach for relating growth rates of pathogen effective population size and time-varying variables that may impact the replicative fitness of a pathogen. The model is applied to real data from rabies virus and *Staphylococcus aureus* epidemics. We find a close correspondence between the estimated growth rates of a lineage of methicillin-resistant *S. aureus* and population-level prescription rates of *β*-lactam antibiotics. The new models are implemented in an open source R package called *skygrowth* which is available at https://mrc-ide.github.io/skygrowth/.

---

Non-parametric population genetic modeling has emerged as a simple, flexible, popular and powerful tool for interrogating genetic sequence data to reveal demographic history (Ho and Shapiro 2011). This approach has proved especially useful for analysis of pathogen sequence data to reconstruct epidemic history and such models are increasingly incorporated into surveillance systems for infectious diseases (Volz et al. 2013). The most commonly used techniques are derivatives of the original *skyline* coalescent model, which describes the evolution of effective population size as a piecewise constant function of time (Pybus et al. 2000). The basic *skyline* model is prone to overfitting and estimating drastic fluctuations in effective population size, so that numerous approaches were subsequently developed for smoothing population size trajectories. Initial approaches to smoothing *skyline* estimators were based on aggregating adjacent coalescent intervals within a maximum likelihood framework (Strimmer and Pybus 2001). Subsequent development has largely focused on Bayesian approaches where a more complex stochastic diffusion process provides a prior for the evolution of a piecewise-constant function of effective population size (Drummond et al. 2005). Non-parametric Bayesian approaches are now the most popular approach for phylodynamic inference and such approaches have illuminated the epidemic history of numerous pathogens in humans and animals (Ho and Shapiro 2011).

To date, all Bayesian non-parametric models have assumed that the effective population size (or its logarithm) follows a stationary stochastic process such as a Brownian motion (Minin et al. 2008; Palacios and Minin 2013). The choice of a stationary process as prior can have large influence on size estimates especially when genealogical data is sparse and uninformative. Genealogies often provide very little information about effective population size near the present (or most recent sample), especially in exponentially increasing populations (de Silva et al. 2012). In such cases, *skyline* estimators with Brownian motion priors on the effective population size may produce estimates which stabilize at a constant level even when the true size is increasing or decreasing exponentially. We argue that in many situations, a more realistic prior can be defined in terms of the growth rate of the effective population size. Below, we describe such a prior based on a simple autoregressive stochastic process defined on the growth rate of effective population size.We show how this prior can lead to substantially different estimates and argue that these estimates are more accurate in many situations. When genealogical data is sparse, our model will retain the growth rate learned from other parts of the genealogy and will correctly capture trends of exponential growth or decline. Even though our approach is non-parametric, we consider its relationship with parametric models of epidemic population genetics to show that our estimates of growth rates of pathogen effective population size are often likely to correspond to growth rates of an infectious disease epidemic.

Smoothing effective population size trajectories using a prior on growth rates also has important advantages when incorporating non-genetic covariate data into phylodynamic inference (Baele et al. 2016). Recent work has focused on refining effective population size estimates using both the times of sequencing sampling (Karcher et al. 2016) or using environmental data which are expected to correlate with size estimates, such as independent epidemic size estimates based on non-genetic data (Gill et al. 2016). Existing statistical models have assumed that the effective population size has a linear or log-linear relationship with temporal covariates. However in many cases, a more realistic model would specify that the growth rate of effective population size is correlated with covariates, as when for example an environmental variable impacts the replicative fitness of a pathogen. We provide a similar extension of previous *skyride* models with covariate data (Gill et al. 2016) to show how such data can be used to test hypotheses concerning their effect and, when a significant effect exists, to refine estimates of both the growth rates and the effective population sizes.

We illustrate the potential advantages of our growth rate model using a rabies virus dataset that has been thoroughly studied using previous phylodynamic methods (Biek et al. 2007; Gill et al. 2016). In particular, we show how our model correctly estimates a recent decline in epidemic size whereas previous models mistakenly predict a stabilisation of the epidemic prevalence. We also apply our methodology to a genomic dataset of methicilin-resistant *Staphylococcus aureus* that had not formally been analysed using phylodynamic methods (Uhlemann et al. 2014). We show how time series on prescription rates of *β*-lactam antibiotics correlate strongly with growth and decline of the effective population size, revealing the impact of antibiotic use on the emergence and spread of resistant bacterial pathogens.

## Methods and Materials

We model effective population size through time as a first order autoregressive stochastic process on the growth rate. This provides an intuitive link between the growth rate of effective population size of pathogens and epidemic size as well as the reproduction number of the epidemic. We further show how to incorporate time-varying environmental covariates into phylodynamic inference.

### Previous Bayesian non-parametric phylodynamic models

Several non-parametric phylodynamic models have been proposed based on Brownian motion (BM) processes and the Kingman coalescent genealogical model (Kingman 1982). In particular, the Bayesian non-parametric *skyride* model uses a BM prior to smooth trajectories of the logarithm of the effective population size (Minin et al. 2008). Let *γ*(*t*) = log(Ne(*t*)) denote the logarithm of the effective population size as a function of time. The BM prior is defined as:

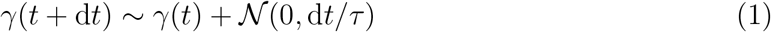

where *τ* is an estimated precision parameter, for which an uninformative Gamma prior is typically used.

This BM prior has been adapted and applied in a variety of ways to enable statistical inference. In the *skygrid* model (Gill et al. 2012), time is discretized and *γ* is defined to be a piecewise constant function of time over a grid with time increments *h*, and the value *γ*_*i*_ is estimated for each interval *i*. Time intervals do not in general correspond to coalescent times in the genealogy. In this case, the BM prior is computed over increments of *γ*:

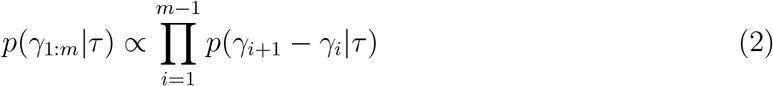

where

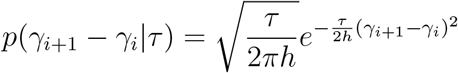

The genealogical data takes the form 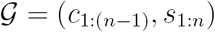 where *c* and *s* are respectively ordered coalescent times (internal nodes of the genealogy) and sampling times (terminal nodes of the genealogy). In the coalescent framework, the sampling times are usually considered to be fixed, so that 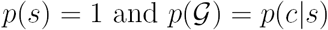. Alternatively, in some variations of this model, a prior *p*(*s*|Ne) is also provided for the sequence of sampling times, making this approach similar to but more flexible than sampling-birth-death-models (Karcher et al. 2016; Volz and Frost 2014).

Given a genealogy, the posterior distribution of the parameters τ and *γ*_1:*m*_ is decomposed as:

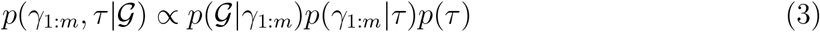

The second term is given by Equation 2 and the last term by the prior on τ. To assist with the definition of the first term, we first denote *A*(*t*) to be the number of extant lineages at time *t*:

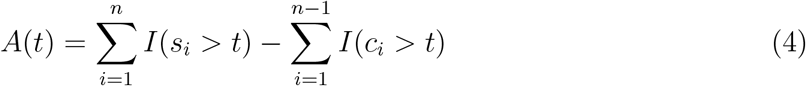

where *I*(*x*) is an indicator function equal to one when *x* is true and equal to zero otherwise. The probability density of the genealogical data given the population size history *γ*_1:*m*_ is then equal to (Griffiths and Tavare 1994):

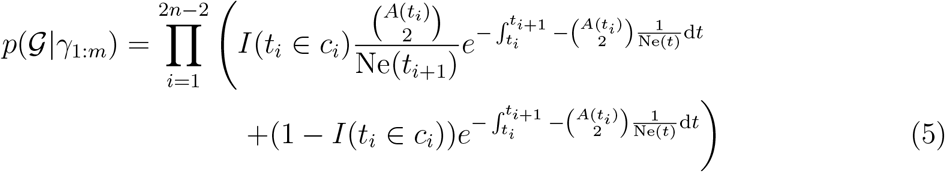

where 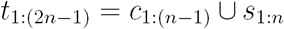 is the set union of sample and coalescent times in descending order.

### Relationship between the growth rate of effective population size and epidemic properties

Several recent studies have investigated the relationship between the effective population size of a pathogen and the number of infected hosts (Koelle et al. 2011; Dearlove and Wilson 2013; Rosenberg and Nordborg 2002). A simple link between these quantities does not exist, since the relationship depends on how incidence and epidemic size change through time (Volz et al. 2009), population structure (Volz 2012), and complex evolution of the pathogen within hosts (Didelot et al. 2016; Volz et al. 2017). Under idealized situations, there is however a simple relationship between the growth rate of effective population size and the growth rate of an epidemic (Frost and Volz 2010; Volz et al. 2013).

Let *Y* (*t*) and *β*(*t*) denote the number of infected hosts and per-capita transmission rate, respectively, as functions of time. Note that *β*(*t*) may depend on the density of susceptible individuals in the population, as in the common susceptible-infected-removed (SIR) model, in which case *β*(*t*) ∝ *S*(*t*)/*N* (Allen 2008). The coalescent rate for an infectious disease epidemic was previously derived under the assumption that within-host effective population size is negligible and that super-infection does not occur (Volz et al. 2009; Frost and Volz 2010):

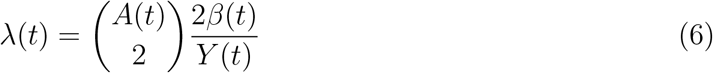

Equating this rate with the coalescent rate under the coalescent model 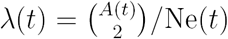 (Kingman 1982) yields the following formula for the effective population size:

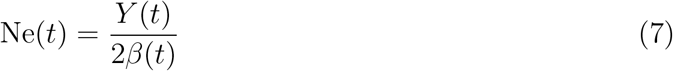

Differentiating with respect to time (denoting with a dot superscript) yields:

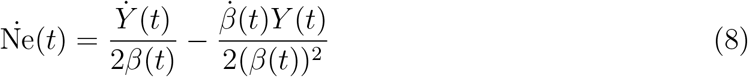

Note that in general the growth rate of the effective population size does not correspond to the growth rate of *Y*, however if the per-capita transmission rate is constant 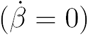, we have 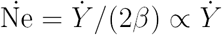. Thus, we expect that over phases of the epidemic where per-capita transmission rates are nearly constant there will be close correspondence between the growth or decline of the effective population size and the growth or decline of the unobserved number of infected hosts. This condition is often satisfied near the beginning of an outbreak which has an exponential phase. It is also often satisfied towards the end of epidemics when the epidemic size is decreasing at a constant exponential rate.

The basic reproduction number *R*_0_ describes the expected number of transmission events caused by a single infected individual in an otherwise susceptible population. By extension, we can define *R*(*t*) as the expected number of transmissions by an infected host infected at time *t* (Fraser 2007). Assuming that all infected individuals are equally infectious (as is the case for example in the SIR model), we have that during periods when the epidemic growth rate is constant, each infected individual transmits at rate *β*(*t*) = *R*(*t*)/*ψ* where *ψ* is the mean duration of infections. With these definitions, the number of infections *Y*(*t*) varies according to the following differential equation:

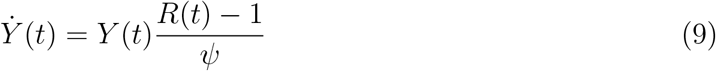

Combining Equations 7, 8 and 9 leads to the following approximate estimator for the reproduction number through time:

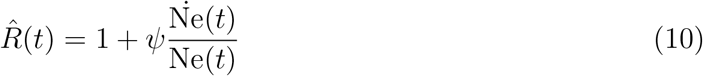

This estimator makes use of the quantity 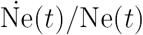 which will be estimated in our model below. Equation 10 is likely to be a good estimator over periods of the epidemic where per-capita transmission rates are invariant. A special case of this occurs at the start of an epidemic, in which case Equation 10 can be used to estimate the basic reproduction number *R*_0_, as previously noted (Pybus 2001).

### A growth rate prior for effective population size

We propose a model in which the growth rate of the effective population size is an autoregressive process with stationary increments. This growth rate is defined as:

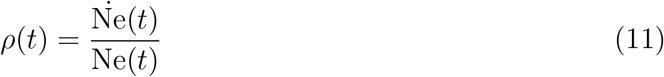

Note that *ρ*(*t*) is a real-valued quantity, with negative and positive values respectively indicating an increase and decrease in the effective population size. In particular, if the population is exponentially growing or declining from *t* = 0 then we have Ne(*t*) = Ne(0)exp(*ρt*) so that *ρ*(*t*) = *ρ* at every time *t* ≥ 0. More generally, we model *ρ*(*t*) using a BM process: *ρ*(*t*) ~ BM(*τ*) (cf Equation 1). To facilitate statistical inference, we work with a discretized time axis with *m* intervals of length *h* as in the *skygrid* model (Gill et al. 2013). We define the growth rate in time interval *i* as:

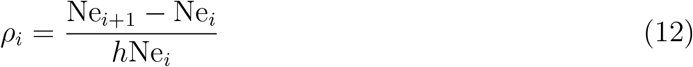

We use the following approximate model for *p*(*ρ*_*i*_+1|*ρ*_*i*_):

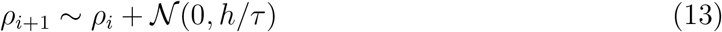

Note that Equation 12 implies *ρ*_*i*_ ∈ (-1/*h*, ∞) since Ne cannot decline below zero, whereas the approximate model in Equation 13 assumes support on the entire real line. We have found performance with this approximate model to be superior to exact models on the log transformation of Ne provided that *h* is small.

With the above definitions, the prior density of a sequence *ρ*_1:*m*_ is defined in terms of the increments:

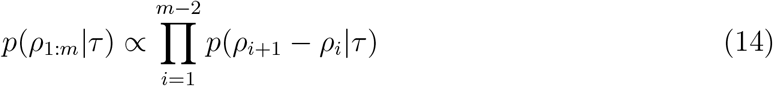

where

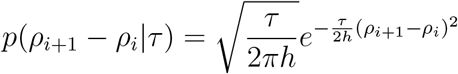

This equation can be compared with the *skygrid* density, Equation 2.

### Incorporating covariates into phylodynamic inference

A simple model was recently proposed for incorporating time-varying covariates into phylodynamic inference with *skygrid* models (Gill et al. 2016). Suppose we observe *q* covariates at *m* time points denoted *X* = (*X*_1:*m*,1:*q*_), and such that observation times correspond to the grid used in the phylodynamic model. The following linear model for the marginal distribution of *γ* with covariate vector *α*_1:*q*_ was proposed:

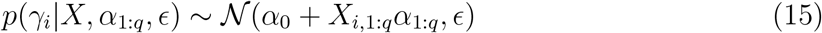

where *α*_0_ is the expected mean of *γ* without covariate effects.

This implies, along with the BM model, the following marginal distribution of the increments:

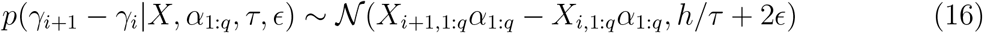

When covariates are likely to be associated with growth rates of the effective population size instead of the logarithm of the effective population size, we can analogously define the density of increments of *ρ*:

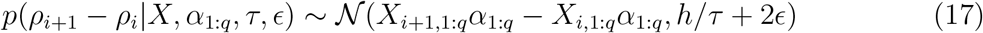

When fitting this model, we drop ∈ for simplicity (as in Gill et al. 2016), and estimate a single variance parameter *τ*.

### Inference and software implementation

Our growth rate model is implemented in an open-source R package called *skygrowth*, available from https://mrc-ide.github.io/skygrowth/, and which includes both maximum a posteriori (MAP) and Bayesian Markov Chain Monte Carlo (MCMC) methods for model fitting.

The MCMC procedure uses a Gibbs-within-Metropolis algorithm that alternates between sampling the growth rate vector *ρ*_1:*m*_ and sampling of the precision parameter *τ*. Metropolis-Hastings sampling is also performed for regression coefficients *α*_1:*q*_ if covariate data is provided with univariate normal proposals. The elements of *ρ*_1:*m*_ are sampled in sequence (from past to present), and multiple Gibbs iterations (by default one hundred) are performed before updating other parameters using Metropolis-Hastings steps.

Maximum a posteriori (MAP) is used as a starting point for the MCMC. The MAP estimator alternates between optimisation of *γ*_1:*m*_ using gradient descent (*BFGS* in R, Goldfarb 1970) and univariate optimisation of *τ* until convergence in the posterior is observed. Approximate credible intervals are provided for the MAP estimator based on curvature of the posterior around the optimum.

## Results

### Simulations

We evaluated the ability of the *skygrowth* model to infer epidemic trends by simulating partially-sampled genealogies from a stochastic individual-based susceptible-infected-recovered (SIR) model. Simulated data were generated using the BEAST2 package MASTER (Vaughan and Drummond 2013), and code to reproduce simulated results is available at https://github.com/emvolz/skygrowth-experiments. The *skygrowth* model was also compared to *skygrid* model as implemented in the *phylodyn* R package (Karcher et al. 2016, 2017) which estimates effective population size using a fast approximate Bayesian non-parametric reconstruction (BNPR). The SIR model was density dependent with a reaction rate *βS*(*t*)*I*(*t*) of generating new infections. Figure 1 shows results of a single simulation with *R*_0_ = 1.3 and 10,000 initial susceptible individuals. Additional simulations are shown in supporting Figure S1. Estimates with *skygrowth* were obtained using the MCMC algorithm and an Exponential(0.1) prior on the precision parameter. We report the posterior means from both *skygrowth* and *skygrid* BNPR. Genealogies were reconstructed by samping 200 or 1000 infected individuals at random from the entire history of the epidemic. In this scenario, both the *skygrowth* and *skygrid* models reproduce the true epidemic trend, capturing both the rate of initial exponential increase, the time of peak prevalence, and the rate of epidemic decline. However, when sampling only 200 lineages (Figure 1A), the genealogy contains relatively little information about later epidemic dynamics, and the *skygrid* estimates revert to a stationary prior producing an unrealistic levelling-off of Ne. Estimates using the *skygrid* BNPR model were highly similar to results using an exact MCMC algorithm for sampling the posterior also included in the *phylodyn* package.

While the results in Figure 1A and B suggest that Ne(t) can serve as a very effective proxy for epidemic size, the degree of correspondence will depend on details of the epidemic model as discussed in the Methods section. Figure 1C and supporting Figure S2 shows a scenario where estimates of *N*_*e*_(*t*) capture the initial rate of exponential growth but fail to estimate the time of peak epidemic prevalence, and the *skygrid* model also fails to detect that the epidemic ever decreases. This scenario was based on a higher *R*_0_ = 5 and only 2,000 initially susceptible individuals, such that almost all hosts are eventually infected and the rate of epidemic decline predominantly reflects the host recovery rate. This is easily understood using the formula Ne(*t*) ∝ *I*(*t*)/*S*(*t*) (cf. Equation 7). When *R*_0_ is large, *S*(*t*) will change drastically over the course of the epidemic. In the later stages, almost all hosts have been infected so that 1/*S*(*t*) is large, producing correspondingly large effective population sizes.

**Figure 1:**
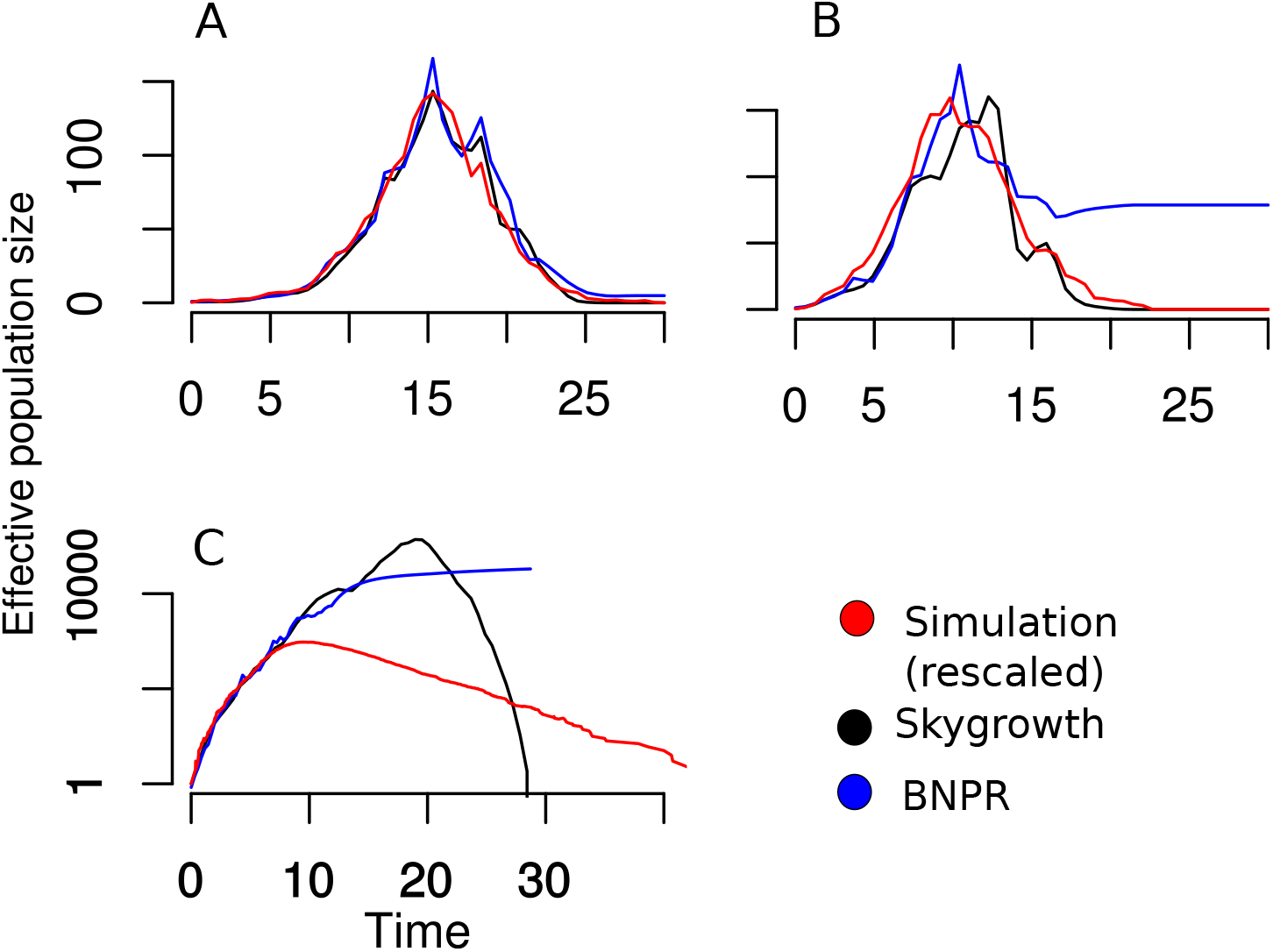
Comparison of effective population size estimates using the *skygrowth* and *skygrid* models applied to data from a susceptible-infected-recovered simulated epidemic. Effective population size estimates are also compared to the number of infected hosts through time under a linear rescaling (red). A. Estimates using a SIR model and simulated genealogy with 1000 sampled lineages and *R*_0_ = 1.3. B. Estimates using a SIR model and simulated genealogy with 200 sampled lineages and *R*_0_ = 1.3. C. Estimates using a SIR model and simulated genealogy with 200 sampled lineages and *R*_0_ = 5.

### Rabies virus

An epidemic of rabies broke out in the late 1970s in the North American raccoon population, following the emergence of a host-adapted variant of the virus called RRV. By the end of the 1990s, this outbreak had spread to a vast geographical area including all Northeast and mid-Atlantic US states (Childs et al. 2000). A sample of 47 RRV isolates has been sequenced in a previous study (Biek et al. 2007), and BEAST (Drummond et al. 2012) was used to reconstruct a dated phylogenetic tree. A standard skyline analysis (Drummond et al. 2005) was performed, which visually suggested a correlation between the inferred effective population size (Ne) and the monthly area newly affected by RRV (hereafter denoted V), but without attempting to quantify the strength or significance of this association.

This data was recently reanalysed using the *skygrid* model with covariates (Gill et al. 2016). No significant association was found between Ne and V, but the authors noted that since V is the newly affected area, V would be expected to be associated with a change in Ne rather than Ne itself. Since the *skyride* method is focused on Ne, like all previous phylodynamic methods, the authors considered the cumulative distribution of V and showed that this is slightly associated with Ne (with a 95% credible interval of [0.18-2.86] on the covariate effect size, Gill et al. 2016). However, this approach is not fully satisfactory. In particular, since V is always positive, the cumulative distribution of V is always increasing, whereas Ne is in principle equally likely to increase or decrease over time. Furthermore both V and its cumulative distribution were considered on a logarithm scale, so that the latter flattens over time by definition.

A more natural solution is to keep the covariate V untransformed, and investigate its association with the growth rate *ρ*(*t*) rather than Ne(*t*) as implemented in our methodology (Figure 2). For this analysis we used exactly the same dated phylogeny as previously published (Biek et al. 2007) (reproduced in Supporting Figure S3). When the covariate was not used (red results in Figure 2), the growth rate was inferred to be positive but declining progressively to zero from 1973 to ~1983, then stable around zero up to ~1990, followed by a period of positive growth until ~2000, after which the growth rate decreased below zero. This implies that the effective population size increased from 1973 to ~1983, then was stable until ~1990, increased to a peak in ~1997 and afterwards decreased. Two waves of spread have therefore been inferred as in previous analyses (Biek et al. 2007; Gill et al. 2016), with the first one starting in the 1970s and ending in ~1983 and the second one lasting from ~1990 to ~1997.

Unfortunately the covariate data V starts in September 1978 and therefore does not cover the first wave. However, the covariate data shows that the epidemic was spreading very quickly between 1992 and 1997, much faster than before or after these dates, and this timing corresponds fairly precisely to the second wave of spread. When the covariate data was integrated into phylodynamic inference, the covariate effect size was found to be statistically significant but only slightly so, with a large 95% credible interval for the covariate effect size of [0.03-4.61] and posterior mean of 1.09. The reconstructed growth rate and effective population size when using the covariate data (blue results in Figure 2) were compatible with results without covariate data. Using additional informative data tightens the credible interval as would be expected, except in the second wave during which the covariate data suggests higher values for both the growth rate and effective population size. The mean posterior growth rate reached a value of about 2.5 per year in the 1990s (Figure 2) and the average generation time of raccoon rabies has previously been estimated to be around 2 months (Biek et al. 2007). We can use Equation 10 to infer a reproduction number of *R* = 1.4, slightly higher than a previous estimate around *R* = 1.1 based on the same data (Biek et al. 2007).

One of the main novel findings of our analysis is that we found a significant decline of the effective population size of raccoon rabies post-2000, whereas previous phylodynamic studies based on the same data found this to be constant (Biek et al. 2007; Gill et al. 2016). Previous methods consider a Brownian motion on the logarithm of Ne, which results in a strong prior that Ne is constant in recent time. By contrast, our model results in the growth rate being a-priori constant, so that the clear decline in growth rate started in the mid-1990s is likely to have continued to the point that the growth rate became negative and Ne declined. Our result is in good agreement with CDC surveillance that shows a clear decline in rabid raccoons after the peak in the mid-1990s (Monroe et al. 2016).

**Figure 2:**
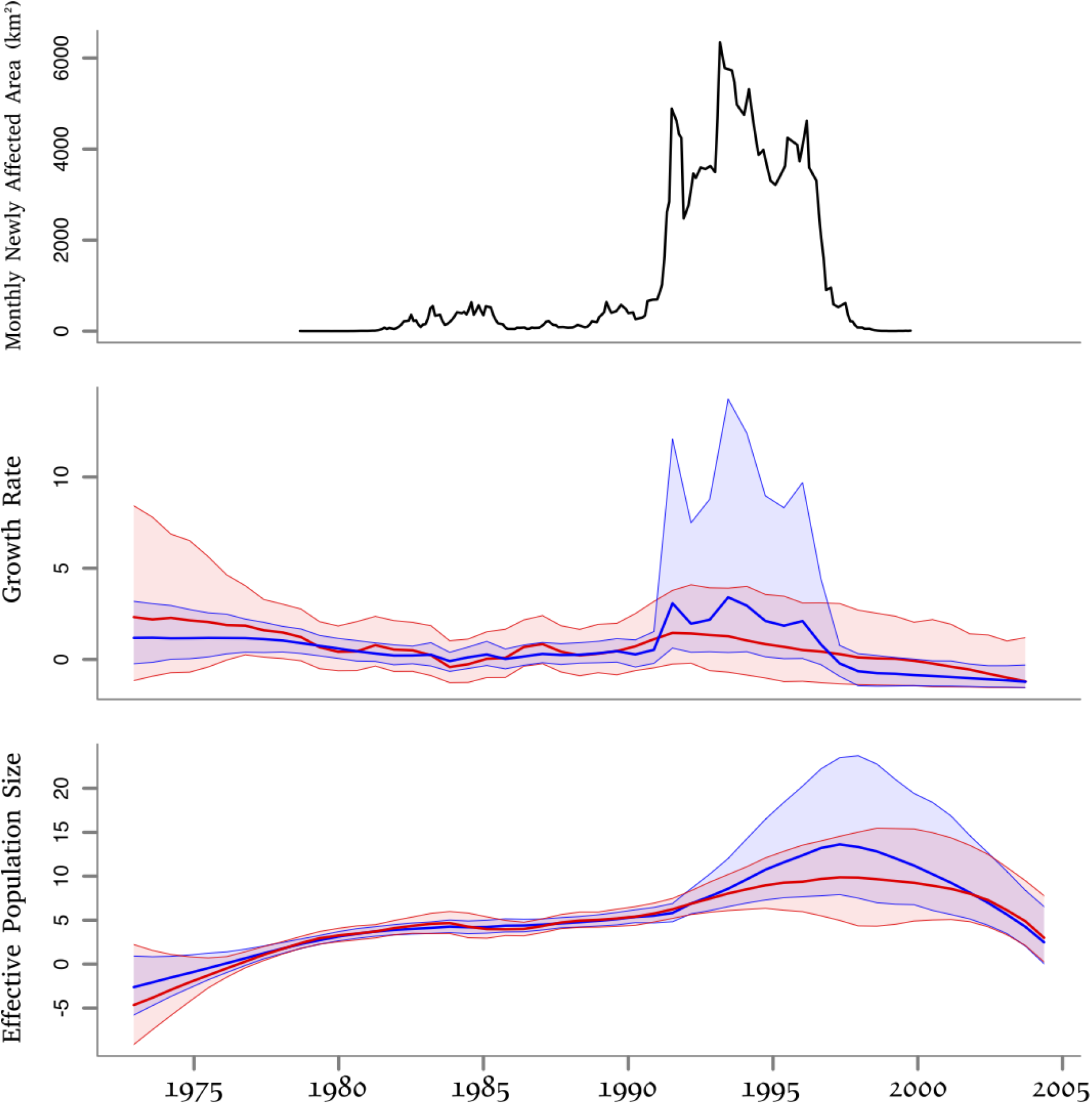
Results on the rabies application. Top: covariate data, representing the area in km^2^ newly affected by rabies recorded monthly between September 1978 and October 1999. Middle: growth rate estimates. Bottom: log effective population size estimates. The middle and bottom plots show results without (red) and with (blue) the use of the covariate data, and with a solid line indicating posterior means and shaded areas indicating the 95% credible regions.

### Staphylococcus aureus USA300

*Staphylococcus aureus* is a bacterium that causes infections ranging from mild skin infections to life-threatening septicaemia. In the 1980s and 1990s, several variants of *S. aureus* have emerged that are resistant to methicilin and other *β*-lactam antibiotics, and collectively called methicilin-resistant *S. aureus* (MRSA) (Chambers and Deleo 2009). MRSA are well known as a leading cause of hospital infections worldwide, but the MRSA variant called USA300 differs from most others by causing infections mostly in communities rather than hospitals. USA300 was first reported in 2000, and has since spread throughout the USA and internationally (Tenover and Goering 2009). A recent study sequenced the genomes from 387 isolates of USA300 sampled from New York between 2009 and 2011, and reconstructed phylogeographic spread that frequently involved transmission within households (Uhlemann et al. 2014).

The USA300 phylogenetic tree (Uhlemann et al. 2014) was dated using a previously described method (Didelot et al. 2012) and a clock rate of ~3 substitutions per year for USA300 (Uhlemann et al. 2014; Alam et al. 2015). We analysed the resulting dated phylogeny (Supporting Figure S4) using our phylodynamic methodology (Figure 3). We initially performed this analysis without the use of any covariate data (red results in Figure 3) and found that the growth rate had been around zero up until 1985, after which it steadily increased until ~1995, and subsequently decreased almost linearly, becoming negative in ~2002 and continuing to decrease afterwards. The effective population size was accordingly found to have been very small until the mid-1990s, to have peaked in ~2002 and to have declined since. These results are in very good agreement with a phylodynamic analysis of USA300 performed using a traditional *skyline* plot on a different genomic dataset (Glaser et al. 2016) as well as USA300 incidence trends (Planet 2017). However, the causes for the recent decline in USA300 are still unclear (Planet 2017). Declines in other MRSA lineages were recently described (Ledda et al. 2017) and have been attributed to improved hospital infection control measures, but this does not apply to the community-associated USA300 lineage.

We hypothesized that the dynamics of USA300 may be driven by the consumption of *β*-lactams in the USA, and we therefore gathered data on this from three different sources covering respectively the periods between 1980 and 1992 (McCaig and Hughes 1995), between 1992 and 2000 (McCaig et al. 2003) and between 2000 and 2012 (CDDEP 2017). There was an overlap of one year between the first and second, and between the second and third of these sources, which was used to scale data for consistency between the three sources. Specifically, values from the second source were scaled so that the 2000 value is equal to the one in the third source, and values from the first source were then scaled so that the 1992 value is equal to the one in the second source. The rescaled data is therefore measured as in the third source, namely in standard units of *β*-lactams (ie narrow-spectrum and broad spectrum penicilins plus cephalosporins) consumed per 1000 population in the USA (CDDEP 2017). This data show that the consumption of *β*-lactams almost doubled between 1980 and 1991, and subsequently decreased to reach around 2010 levels comparable to the early 1980s (Figure 3). These trends on *β*-lactams consumption therefore appear to be very similar to the ones observed for the USA300 growth rate without the use of covariates (red results in Figure 3). To confirm this observation, we repeated our phylodynamic analysis with integration of the *β*-lactam use as a covariate (blue results in Figure 3). We found that the covariate was significantly associated with growth rate, with a mean posterior effect of 0.48 and 95% credible interval [0.18-0.71]. The growth rate dynamics inferred when using covariate data was almost identical to those inferred without the use of covariate data, except for a clear reduction of the width of the intervals which reflects the gain in information when combining two independent types of data.

**Figure 3:**
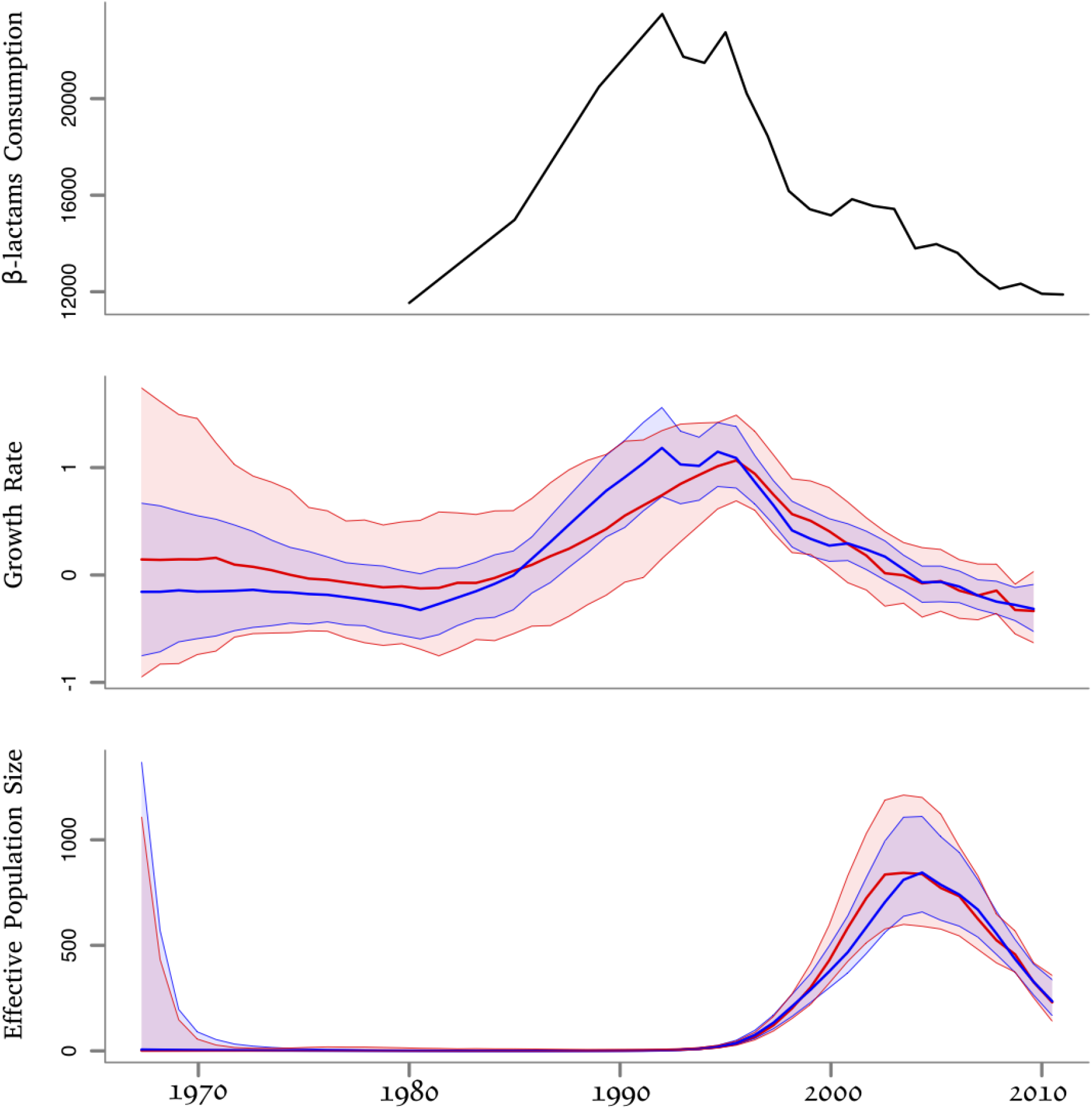
Results on the USA300 application. Top: covariate data, representing the consumption of *β*-lactams between 1980 to 2012 in the USA, measured in standard units per 1000 population. Middle: growth rate estimates. Bottom: log effective population size estimates. The middle and bottom plots show results without (red) and with (blue) the use of the covariate data, and with a solid line indicating posterior means and shaded areas indicating the 95% credible regions.

Our analysis therefore suggests that the rise in *β*-lactams consumption in the 1980s was responsible for the emergence of the highly successful USA300 lineage. From the mid-1990s, the use of *β*-lactams has declined, both due to an overall reduction in antibiotic use and a diversification of the type of antibiotics prescribed (McCaig et al. 2003; CDDEP 2017), and the growth rate of USA300 has consequently decreased. Importantly, the consumption of antibiotics is expected to be associated with the growth rates of resistant bacterial pathogens, rather than with their effective population sizes, which here is not at all correlated with the covariate (Figure 3). Amongst pairs of individuals thought to have infected one another within households, the distribution of genomic distance had a mean of 4 substitutions (Uhlemann et al. 2014), and this represents on average twice the number of substitutions occurring during an infection when accounting for within-host diversity (Didelot et al. 2012, 2014, 2016). Given that the molecular clock rate of USA300 is approximately 3 substitutions per year (Uhlemann et al. 2014; Alam et al. 2015), the average duration of infections in this outbreak is around eight months. In the first half of the 1990s, the growth rate peaked around 1 per year (Figure 3) and using Equation 10 we estimate that the reproduction number was around *R* = 1.6, which is in good agreement with the recent estimate *R* = 1.5 for MRSA in the US population (Hogea et al. 2014). The fact that this estimate is only modestly above the minimum threshold of *R* = 1 required for outbreaks to take place could help explain why the USA300 is declining, even though *β*-lactams are still widely used. The consumption level may have lowered below the threshold caused by the fitness cost of resistance, as previously discussed for other resistant bacteria (Whittles et al. 2017; Dingle et al. 2017).

## Discussion

Many environmental covariates, particularly those with a mechanistic influence on replicative fitness of pathogens, are closely related to the growth rate of epidemic size but not necessarily related to absolute epidemic size. We have found that these relationships can be inferred from random samples of pathogen genetic sequences by relating environmental covariates to the growth rate of the effective population size. This enables the estimation of the fitness effect of environmental covariates as well as the prediction of future epidemic dynamics should conditions change. We have found a clear and highly significant relationship between the growth and decline of community-associated MRSA USA300 and the population-level prescription rates of *β*-lactam antibiotics (Figure 3). This relationship is not apparent when comparing antibiotic usage directly with the effective population size of MRSA USA300. Our methodology focused on growth rate is therefore well suited to investigate the drivers of antibiotic resistance, compared to previous phylodynamic methods focused on the effective population size.

The *skygrowth* model can provide a more realistic prior for many infectious disease epidemics where the growth rate of epidemic size is likely to approach stationarity as opposed to the absolute effective population size. Conventional *skyride* and *skygrid* models are prone to erroneously estimating a stable effective population size when genealogical data is uninformative, as for example when estimating epidemic trends in the latter stages of SIR epidemics (Figure 1). The *skygrowth* model will correctly predict epidemic decline in this situation. Moreover, under ideal conditions, the estimated growth rate can be related to the reproduction number of an epidemic, and the *skygrowth* model provides a simple non-parametric estimator of the reproduction number through time given additional information about the natural history of infection (Equation 10). Caution should be exercised when using the effective population size as a proxy for epidemic size, as the relationship between the two is complex (cf. Simulation results). In general, there will be close correspondence between the growth of epidemic size and growth of effective population size during periods where the growth rate is relatively constant.

The methods presented here can be applied more generally to evaluate the role of antibiotic stewardship, vaccine campaigns, or other public health interventions on epidemic growth rates. Some environmental covariates, such as independent prevalence estimates, may be more closely related to effective population size rather than growth rates, and future work is indicated on the development of regression models in terms of both statistics. More complex stochastic models can also be considered, such as processes with both autoregressive and moving average components. A variety of mathematical models have been developed to explain de novo evolution of antimicrobial resistance as a function of population-level antimicrobial usage (Bonhoeffer et al. 1997; Austin et al. 1999; Spicknall et al. 2013; Whittles et al. 2017), and an important direction for future work will be the development of parametric and semi-parametric structured coalescent models (Volz 2012) that can be applied to bacterial phylogenies featuring a mixture of antibiotic sensitive and resistant lineages. This methodology will allow us to estimate key evolutionary parameters, such as the fitness cost and benefit of resistance, or the rate of mutation from sensitive to resistant status, which are needed to make well informed recommendations on resistance control strategies.

